# Maternal Alcohol Drinking Patterns Predict Offspring Neurobehavioral Outcomes

**DOI:** 10.1101/2024.03.06.583121

**Authors:** Abbey Myrick, Diane Jimenez, Belkis Jacquez, Melody S. Sun, Shahani Noor, Erin D. Milligan, C. Fernando Valenzuela, David N. Linsenbardt

## Abstract

**SIGNIFICANCE:** The timing, rate, and quantity of gestational alcohol consumption, collectively referred to here as Maternal Drinking Patterns (MDPs), are of known importance to fetal developmental outcomes. Though studies in rodents exist that have investigated the impact of gestational alcohol drinking characteristics, few have sought to determine the impact of MDPs on offspring behavioral outcomes.

**METHODS:** We first used specialized equipment to record the precise amount and timing of binge alcohol consumption in pregnant mouse dams, and then characterized MDPs using Principle Component Analysis (PCA). We focused these analyses on the first fifteen minutes of every gestational drinking session when dams consumed the majority of each session’s alcohol (a phenomenon known as front-loading), as well as the entire 2 hour session across all days of gestation. We next tested offspring in open field and rotarod assays and evaluated these behavioral results in the context of MDPs.

**RESULTS:** Male alcohol exposed mice exhibited longer latencies to fall on the rotarod compared to their controls, which we attribute to a delayed decrease in body weight-gain not observed in females. This effect was found to be associated with MDPs within the first fifteen minutes of drinking, but not other MDPs. Female alcohol exposed mice had significantly reduced total locomotor activity in the open field compared to controls, and this effect was also associated with MDPs but only of the entire drinking session. Surprisingly, total gestational alcohol consumption alone was not associated with any particular behavioral outcome. Furthermore, we replicated robust behavioral data demonstrating development of allodynia in alcohol exposed mice where it did not develop in controls.

**CONCLUSIONS:** To our knowledge, this report represents the highest resolution assessment of alcohol drinking throughout gestation, and one of few to have identified relationships between specific alcohol MDPs and neurobehavioral outcomes in offspring. Specifically, based on characteristics of the PCA groups, we found evidence that the rate of alcohol front-loading leads to developmental delays in males, whereas an interaction of front-loading rate and duration, overall persistence, and total amount consumed lead to a female-only decrease in locomotor activity. Beyond these results, we provide a method for precise and accessible tracking of such data.

## Introduction

Alcohol (ETOH) is a known teratogen, with fetal exposure leading to a range of deleterious neurobehavioral outcomes, including learning and memory deficits^1,2^, impaired motor coordination^3^, increased anxiety^4^, and greater risk for developing touch hypersensitivity, a peripheral neuropathy referred to as allodynia pain^5^. These outcomes are due in large part to the broad impacts of prenatal alcohol exposure (PAE) on neural development, which are diagnosed in the clinic as Fetal Alcohol Spectrum Disorders (FASDs). FASDs affect as many as 5% of the United States population, making it the most prevalent, modifiable neurological disorder^6^. Unfortunately, ETOH consumption during pregnancy continues, and there are recent data showing it may be increasing^7,8^.

While the highest rates of ETOH consumption occur during the first trimester, often before women are aware they are pregnant, its use is also common at later stages of pregnancy including the 2^nd^ and even 3^rd^ trimester^9^. This is an important consideration, because gestational timing of alcohol exposure has been shown to have differential impacts on neurodevelopmental outcomes, presumably due to the developmental stage of the fetus. For example, Ikonomidou et al (2020) found region-specific differences in neurodegeneration as a result of specific gestational exposure windows^10^. Furthermore, Lipinski et al (2012) demonstrated differences in facial dysmorphology in mice exposed to alcohol on GD7 versus GD8.5, with parallel changes to brain mass^11^. While studies such as these emphasize the importance of explicitly evaluating the impact of gestational alcohol exposure timing to neurobehavioral outcomes of offspring, few studies to date have done so.

Another critically important maternal alcohol drinking characteristic is the rate of consumption. Rapid alcohol consumption, or “binge-drinking”, has been shown to have a more detrimental outcome to neurodevelopment than quantity of alcohol alone^12,13^. For example, Bonthius et al (1990) exposed developing rats using a larger alcohol dose infused slowly (6.6 g/kg), or a lower dose (4.5 g/kg) infused more quickly, and found that the lower dose led to higher blood ethanol concentrations (BECs) and greater cell loss in the hippocampus and cerebellum versus larger dose. Thus, the timing, amount, and rate of maternal alcohol consumption, which we collectively refer to as maternal drinking patterns (MDPs), are critically important factors that should be evaluated in PAE studies.

Investigating the consequences of MDPs on PAE individuals in the clinic is challenging, largely because it is often impossible to obtain precise records of amount and timing of exposure. Preclinical studies circumvent this issue, but until recently there was a lack of practical and efficient methods for monitoring MDPs accurately. To this end, we used a Volumetric Drinking Monitor (VDM) hardware system together with the drinking-in-the-dark (DID) binge alcohol drinking assay to extremely accurately capture MDPs. We subsequently evaluated the behavior of these dam offspring, and for the first time identified MDPs associated with sex-specific neurobehavioral outcomes. Specifically, we found MDPs associated with offspring performance on body weight gain and the rotarod ataxia assay in males, and ambulatory activity in a novel open field assay in females. These findings highlight the multifaceted contributions of MDPs to fetal neurobehavioral outcomes, and in turn the importance of considering them in PAE studies.

## Methods

### Animals

40 8-week old male and female C57BL/6J mice were obtained from The Jackson Laboratories (JAX) to be used as breeders for the production of prenatally alcohol exposed and control subjects. Upon arrival, mice were single housed in 6 × 10 × 5 inch amber cages with wire top food hoppers and standard water bottles containing tap water. Mice were acclimated to the room for two weeks under a reverse 12-hour light/dark cycle before study initiation. All animals had *ad lib* access to LabDiet 5001, a mouse chow previously shown by our lab to support reliable ETOH consumption^14^. Water was always available except for 2 hours a day in ETOH exposed mice (see below).

### Drinking-in-the-Dark (DID)

For the purposes of precisely evaluating drinking patterns with microliter precision^15^ animals were provided with either 20% ETOH or water via a Volumetric Drinking Monitor system (VDM; Columbus Instruments, Columbus OH) using standard DID procedures; standard water bottles were replaced daily with specialized sippers for 2 hours, 3 hours into the dark cycle. Water for all sipper drinking solutions came from the same source within the New Mexico Animal Resource Facility as used for standard housing water bottles.

### Blood Ethanol Concentrations (BECs)

Blood samples were drawn from the peri-orbital sinus immediately following the conclusion of the 2-hour drinking period on the 15^th^ day following breeding, which will be referred to as the 15^th^ day of gestation (GD15). Blood plasma was isolated and kept frozen at -20ºC until analysis of BECs using an Analox Alcohol Analyzer (Analox Instruments, Lunenburg, MA).

### Breeding/Offspring

Following a two week DID acclimation period, each female was housed with a male immediately after daily sipper removal for three consecutive days for ∼5 hours; females were removed from male cages at least an hour before lights on. If a dam did not become pregnant, she was re-bred two weeks later. Drinking sessions were terminated as soon as a dam gave birth. Offspring were weaned 22 days following birth by placing with littermates into same-sex groups, were housed in a 12-hour light/dark cycle, and were tested throughout development on several assays previously used to evaluate the consequences of prenatal alcohol exposure on behavior detailed below^16^.

### Rotarod

On postnatal days (PD) 53-58 offspring were next tested during the dark cycle under dim red light. Each animal performed 6 total trials; three consecutive trials each day for two days. Each trial consisted of placing the animal one of several walled rods initially rotating 4 revolutions per minute (RPM). After 10 seconds, the speed gradually increased to 20 RPMs. The time at which the animal fell off the rod was recorded as the latency to fall.

On postnatal days (PD) 53-58 offspring were next tested during the dark cycle under dim red light, according to previously published methods^17^. Briefly, each animal performed 6 total trials; three consecutive trials each day for two straight days. Each trial consisted of placing the animal on one of several walled rods initially rotating 4 revolutions per minute (RPM). After 10 seconds, the speed gradually increased to 20 RPMs. The time at which the animal fell off the rod was recorded as the latency to fall.

### Open Field

On PD 71-96 offspring were tested in a novel open field behavioral assay^18^ under ∼150 lumens of white light. A single mouse at a time was placed in the center of a white square (50 × 50 cm) acrylic arena which it was allowed to freely explore for 30 minutes. The arenas were wiped down with 70% isopropanol between each mouse. An overhead Noldus EthoVision camera (Leesburg, VA) tracked the location and distance traveled throughout. Custom MATLAB code was used to quantify time spent in the center of the box versus the outer edges.

### Minor Chronic Constriction Injury

In order to model allodynia, a Minor Chronic Constriction Injury (mCCI)—the well-characterized sciatic nerve chronic constriction injury (CCI) model of peripheral neuropathy was applied^19^ with modifications, as previously reported^5,20^ to create a minor injury, as briefly described here. Following isoflurane anesthesia (1.5-2.0% volume in oxygen, 2.0 L/min), dorsal left thigh was shaved and cleaned using 70% ethanol that was air dried prior to surgery. Using aseptic procedures, a 15-scalpel blade was used to make a 2 cm incision in the skin just caudal to the femur, followed by a blunt dissection to carefully isolate the sciatic nerve by separating the overlying muscular fascia. Sterile plastic probes were then used to find and lift the sciatic nerve out of the muscle pocket, and a single piece of 6-0 Chromic Gut (Ethicon: Cat # 796G) suture was snuggly tied around the sciatic nerve, proximal to the trifurcation, without pinching the nerve. The sciatic nerve and surrounding muscle were kept irrigated throughout the procedure, using isotonic sterile saline (Hospira: Cat # NDC 0409-4888-03) to prevent dehydration. Following ligation, the nerve was gently placed back into position using sterile plastic probes, and overlying muscle fascia was sutured closed using one sterile 4-0 Silk Suture (Ethicon: Cat # 1677G). The overlying skin was closed using two 9mm Reflex™ Wound Clips (Kent Scientific Corp: Cat # INS750344), which were removed 7 days later. Sham surgeries were performed identically to the mCCI surgeries. Procedures were completed within 15-20 min. Mice fully recovered from anesthesia within approximately 10 min and were monitored daily for post-operative complications; body weight, wound condition, hindpaw autotomy, activity, and grooming behaviors were checked to confirm healthy recovery. All mice in this study recovered completely from the surgery, without any abnormalities, and remained in the study.

### Assessment of Allodynia

Mechanical allodynia was assessed applying the von Frey test to identify hindpaw threshold responses adapted for mice^21^. Each experimental condition was represented on each day of von Frey testing. Timepoints for hindpaw threshold assessment were chosen based on prior reports demonstrating the onset, magnitude and duration of allodynia^20^. Mice were habituated to the experimenters and testing environment for 45-60 minutes within the first three hours of lights for four sequential days, followed by baseline (BL) assessment, one/day, separated by at least 48 hours. Hindpaw threshold responses to light mechanical stimuli were assessed using nine calibrated von Frey monofilaments (touch test sensory evaluator: North Coast Medical: Cat # NC12775) applied for a maximum of 3 seconds (a metronome was set to 60 beats per minute) to the plantar surface of both the left (ipsilateral to CCI) and right (contralateral to CCI) hindpaws, with randomized ipsilateral and contralateral testing order to avoid order effects. Inter-trial stimuli intervals were 30 seconds apart. The log intensity of the nine monofilaments used is defined as log_10_ (grams x 10,000) with the range of intensity being as follows reported in log (grams): 2.36 (0.022g), 2.44 (0.028g), 2.83 (0.068g), 3.22 (0.166g), 3.61 (0.407g), 3.84 (0.692g), 4.08 (1.202g), 4.17 (1.479g), and 4.31 (2.042g). Each paw was initially presented with the monofilament marking of 3.22 (0.166 g) (middle of the assay scale) with subsequent monofilaments used based on the response/non-response of the mouse to the previous monofilament tested. No response observed by the stimulus presented from the 3.22 monofilament dictated that the next “greater” monofilament to be applied (e.g., 3.61). If a response was elicited by the stimulus presented from the 3.22 monofilament, the next “lighter” monofilament was tested (e.g., 2.83). This pattern of monofilament stimulus presentations persisted with up to six presentations applied each paw.

### Statistical Analysis

Gestational drinking data and estimated BECs were evaluated using repeated measures ANOVAs with day as the within-subjects factor and fluid type as the between groups factor. Relationship between ETOH consumption and BECs was determined using simple linear regression. Estimated BECs were calculated using the function that described the fit between ETOH and BEC. Three separate principle component analysis (PCAs) were run in MATLAB to cluster variance in dam drinking patterns along PC1. The first PCA focused on dam drinking patterns for GD15 only, when BECs were taken. The second analyzed all days of gestation, and the third looked at all days of gestation but only the first 15 minutes of each drinking session. Offspring were then grouped by respective dam’s PCA cluster (indicated as Group 1 or Group 2) and performance in the open field and rotarod were analyzed using nested T-tests. Each PCA was compared to open field and rotarod data separately. Rotarod and open field data was analyzed using ANOVAs unless otherwise stated. Sex was analyzed separately and any sex-specific effects were reported. Nested t-tests found no significant impact of litter on either rotarod or open field performance. Allodynia data was quantified using the total number of positive and negative responses summed for each monofilament presented, and entered into the computer software program, PsychoFit (http://psych.colorado.edu/~lharvey: RRID: SCR_015381) to determine the absolute withdrawal threshold (50% paw withdrawal threshold) by applying a Gaussian integral psychometric function to determine the maximum-likelihood fitting method, as previously described^22^, and the 50% paw withdrawal thresholds were plotted and used for statistical analysis. 3-way ANOVAs were conducted for males and females separately in Prism to evaluate touch sensitivity across multiple days.

## Results

### Total gestational fluid consumption

The mean fluid consumed throughout gestation can be seen in Figure 1A. Repeated measure ANOVA revealed a significant main effect of fluid type [F (1, 31) = 84.50, p < 0.0001] as well as a significant fluid type by day interaction [F (19, 589) = 1.793, p = 0.0205]; effects driven principally by greater volume consumed in the water assigned control group.

**Figure 1.**
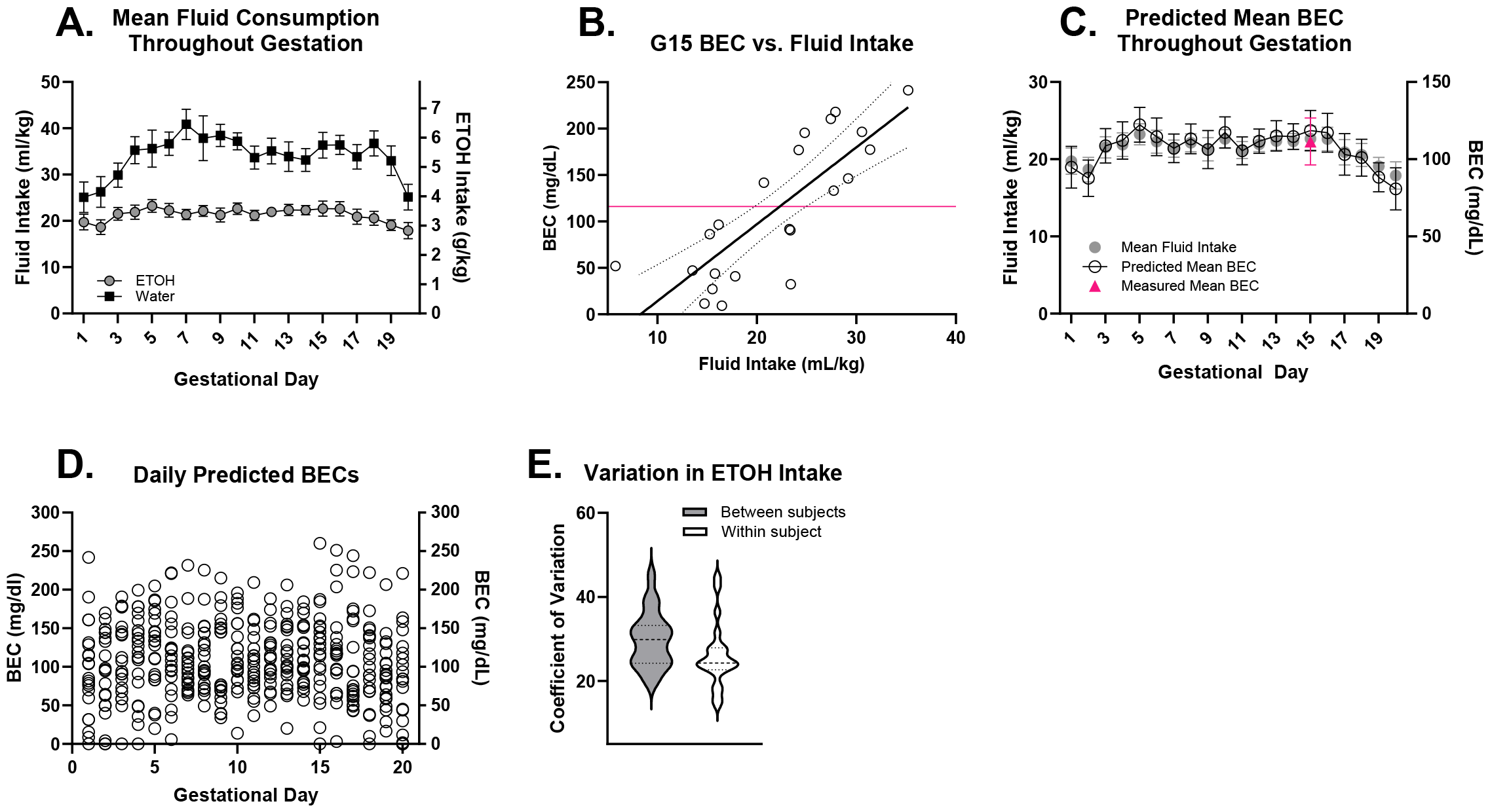
A) Mean fluid intake throughout gestation. Dams assigned water consumed more fluid volume by weight versus ETOH-assigned dams (P<0.0001). B) A significant correlation was found between measured BECs and fluid intake on gestational day 15 (p<0.0001). C) Predicted mean BECs generated using mean fluid intake across all days of gestation. D) Individual predicted BEVs based on daily intake. E) A high degree of variation in predicted BECs is due to variation in ETOH consumption between and within subjects.

### Gestational BECs

Mean BECs collected on the 15^th^ day of gestation can be seen in Figure 1B. As expected, simple linear regression revealed a significant positive relationship between BEC and fluid intake [*R*^2^= 0.6484, F = 36.88, p <0.0001, CI = 0.95]. The function describing this relationship (Y = 8.286*X - 68.68) was then used to generate predicted daily BECs based on mean fluid intake (Figure 1C), which confirmed relatively stable BECs and intake as a mean of treatment group throughout gestation. However, further evaluation of gestational drinking and predicted BECs revealed a high degree of variation (Figure 1D), which was found to be the result of both between- and within-subject variability (Figure 1E).

### Pattern of gestational fluid consumption

On GD15, the frontloading behavior can be seen in Figure 2A, wherein a mixed-effects analysis found a main effect of treatment, where consumption rate is highest within the ETOH group during the first fifteen minutes of the session (Fig. 2B), a characteristic that continued throughout gestation [F(10.20, 339.2) = 2.132, p<0.0001]. Interestingly, a linear regression revealed a significant increase in frontloading among ETOH dams as gestation progressed [(ETOH: F(1, 438) = 18.31, R^2^ = 0.04012, p<0.0001)(Water: F(1, 278) = 0.5671, R^2^ = 0.002036, p = 0.4521)]. To further investigate differences in dam drinking patterns, we began with a PCA of the entire GD15 session (PCA1). Our analysis produced two clusters of variance (Fig. 3A) where Group 1 contained animals that achieved higher BECs [F(11, 9) = 1.161, p = 0.0001] than Group 2 and whose mean drinking patterns can be seen in Figure 3B. We then ran a second PCA (PCA2) for the first fifteen minutes of every session across all days of gestation (Fig. 3C and 3D). Finally, our third PCA (PCA3) leveraged the entire session across all days of gestation (Fig. 3E and 3F). Using the average cumulative drinking across all days of gestation, dams were also divided into high and low drinking categories based on the mean (3.303 g/kg) in order to assess the impact of total ETOH intake on offspring behavior.

**Figure 2.**
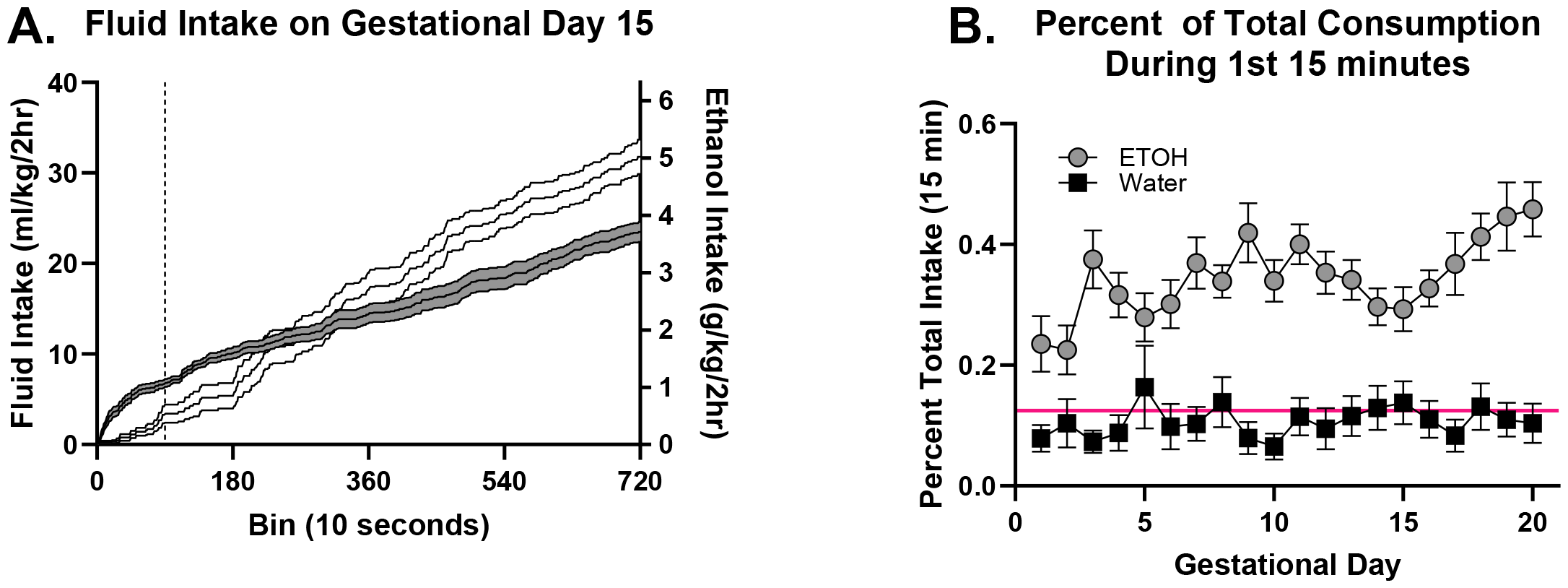
A) Cumulative fluid consumption on gestational day 15. ETOH animals have the highest fluid intake rate during the first 15 minutes (indicated with a dashed line) of the drinking session. B) ETOH exposed dams consume a significantly higher percentage of their total fluid within the first 15 minutes of a drinking session compared to water exposed dams across gestation (p<0.0001).

**Figure 3.**
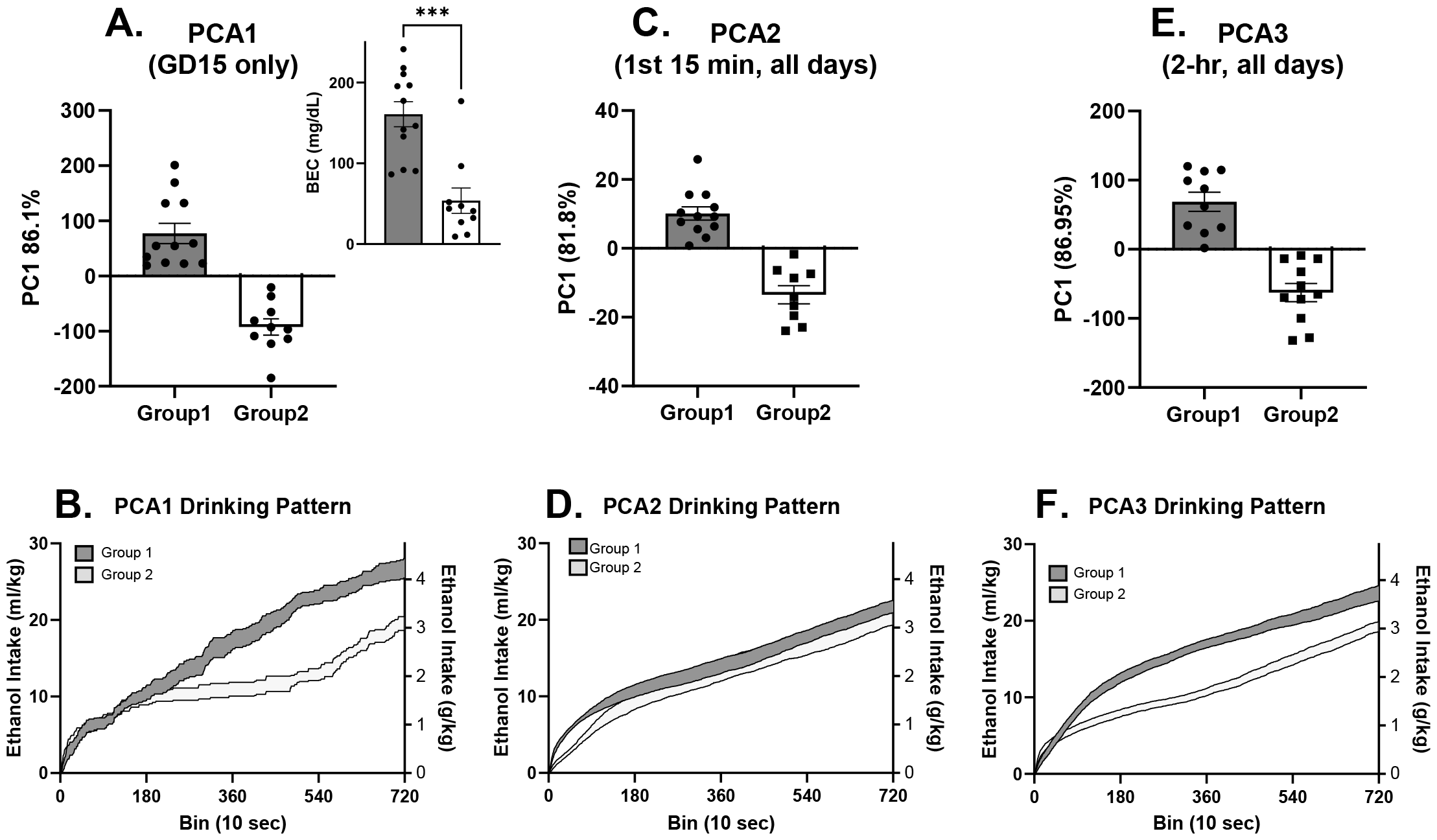
A) PCA on D15 when BECs were taken; group one accounts for 86.1% of all variance, group 2 accounts for 12.45%. Inset indicates comparative BECs, with group 1 exhibiting roughly three times greater BECs on average (P = 0.0001). B) Cumulative drinking pattern as a function of PCA group on GD15. C) PCA clustering of the first fifteen minutes of each session across all days of gestation. D) Mean drinking pattern of PCA2. E) PCA of drinking pattern for whole 2-hour session across all days of gestation. F) Mean drinking pattern of PCA3.

### Offspring body weights

While lower birth weights are a known consequence of PAE^23^, a paired t-test did not indicate significant differences in percent change of gestational dam weight between treatment groups [t (4) = 1.931, p = 0.1256) (Fig. 4A). Mann-Whitney tests revealed no significant treatment differences for the number of pups born to a litter (Fig. 4B) [p = 0.3326, U = 95, n = 12 (water), n = 20 (ETOH)]. Likewise, an unpaired t-test found no differences in average pup weight per litter at postnatal day 10 (Fig. 4C) [t(30) = 0.2779, p = 0.7830]. However, by mean postnatal day 57(SEM±0.23), a two-way ANOVA of body weights identified a significant main effect of treatment [F(1, 87) = 13.80, p = 0.0004], with post-hoc tests confirming that ETOH exposed males were driving the effect (Fig. 4D). [(Males; N1 = 34, N2 = 16, t = 3.616, p = 0.0010)(Females; N1 = 24, N2 = 17, t = 1.681, p = 0.0964)].

**Figure 4.**
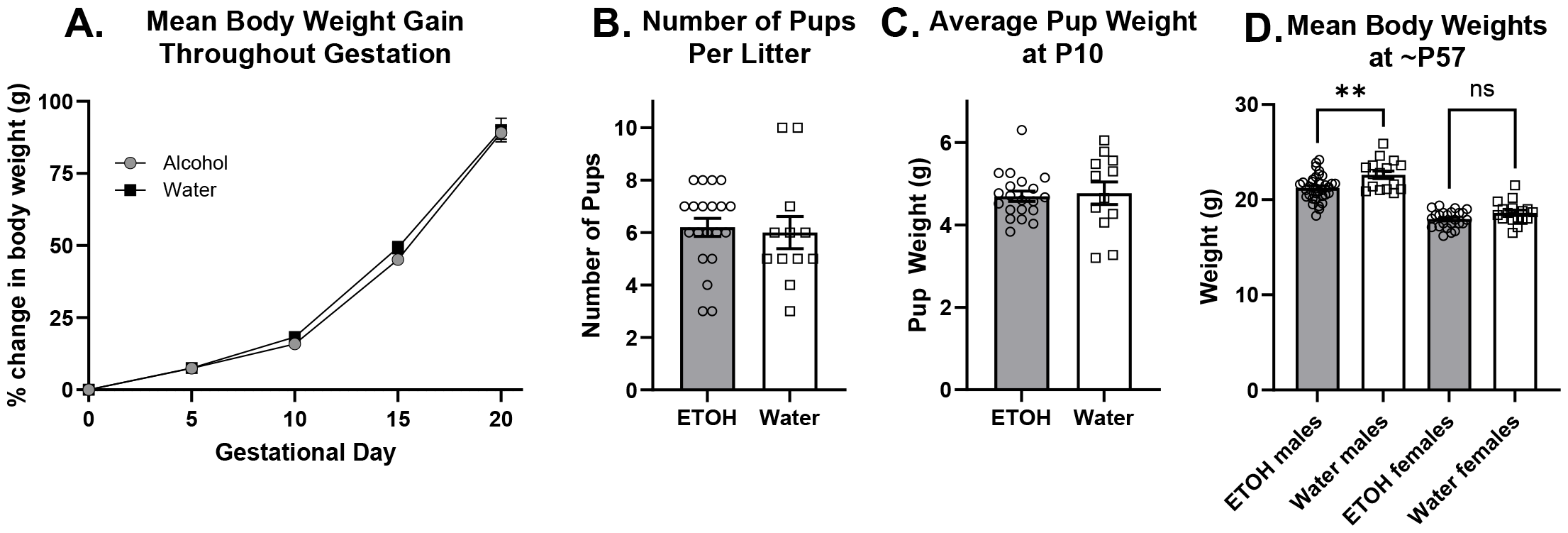
A) Weight gain across gestation was not significantly different between treatment groups. B) No significant differences in number of pups born between treatment groups. C) No significant treatment difference in average pup weight per litter. D) By mean postnatal day 57, a treatment difference in body weights emerged among males (p = 0.0010).

### Rotarod

The results of motor coordination assessment using the accelerating rotarod can be seen in (Fig. 5A). Repeated measure ANOVA conducted on sex separately identified significant main effects of trial in both males (F(3.464, 246.0) = 20.24, p<0.0001) and females (F(4.284, 304.1) = 35.44, p<0.0001], resulting from increases in time spent on the rotarod over trials. A main effect of treatment was found among males [F(1,71) =4.669, p = 0.0341], but no significant treatment difference was present in females [F(1, 71) = 0.01771, p = 0.8945]. This sex dependent treatment effect became more apparent wh en analyzed alongside weight (Fig. 5B), and a simple linear regression found weight could predict rotarod performance (fitted regression model: Y = -9.756*X + 285.5)[F(1, 89) = 20.05, *R*^2^ = 0.1839, p<0.0001).

**Figure 5.**
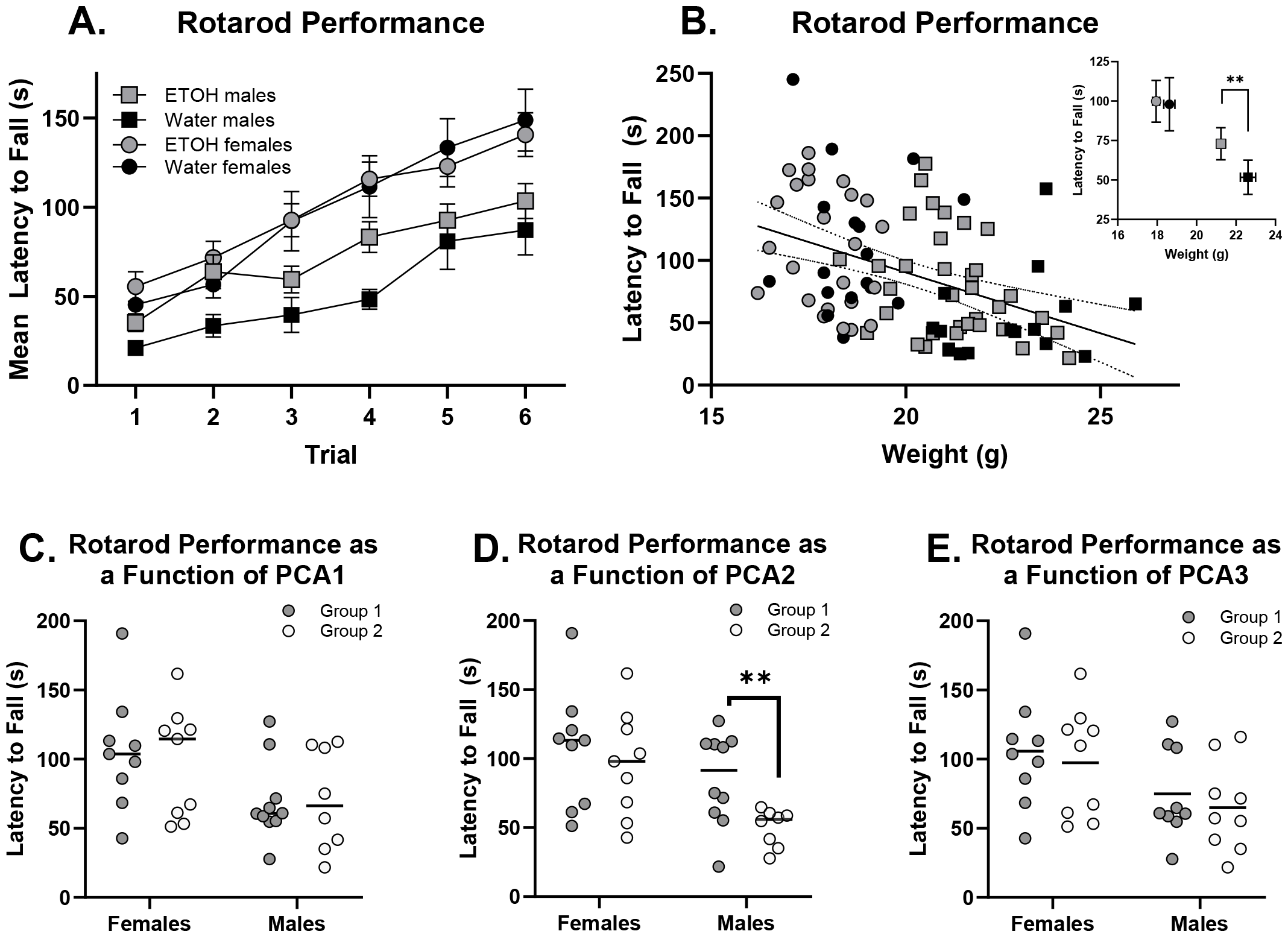
A) A main effect of treatment was found in rotarod performance in male mice (p = 0.0341) but not in female mice (p = 0.8945). B) Weight is predictive of rotarod performance (p<0.0001). C) No correlation with PCA1 and rotarod performance (females; p = 0.8553)(males; p = 0.6838). D) Significant correlation between rotarod performance and dam drinking pattern in PCA2 among males (p = 0.0021) but not females (p =0.4593). E) No correlation was found between PCA3 drinking patterns and rotarod performance (females; p = 0.7972)(males; p = 0.7505).

Performance was then assessed based on dam drinking patterns as defined by PCA grouping. A nested t-test found no significant relationship between offspring performance on the rotarod in either sex in PCA1 [(females; t(16) = 0.1854, p = 0.8553)(males; t(16) = 0.4148, p = 0.6838)] (Fig. 5C). But the same analysis with PCA2 revealed a sex-specific correlation to dam drinking pattern and performance among males [t(16) = 3.665, p = 0.0021] but not females [t(16) = 0.7582, p =0.4593] (Fig. 5D). No relationship to rotarod performance was found with PCA3 among females [t(16) = 0.2614, p = 0.7972] or males [t(16) = 0.3235, p = 0.7505] (Fig. 5E).

No significant relationship between average cumulative maternal alcohol consumption and offspring rotarod performance was found for either sex (Females: p = 0.2369; Males: p = 0.7942, data not shown).

### Open Field

A treatment effect was found in total locomotor activity using an unpaired t-test (Fig. 6A) [t(149) = 2.604, p = 0.0101] and a nested t-test found the females were driving the treatment effect (Fig. 6B) [t(25) = 2.315, p = 0.0291], while the males had no significant treatment difference [t(25) = 1.056, p = 0.3007]. An unpaired t-test found no significant treatment effect when time spent in the center of the box was analyzed as a percent of total time in the box [(both sexes; t(154) = 0.6924, p = 0.4897)(females; t(74) = 1.694, p = 0.0945)] (Fig. 6C and 6D).

**Figure 6.**
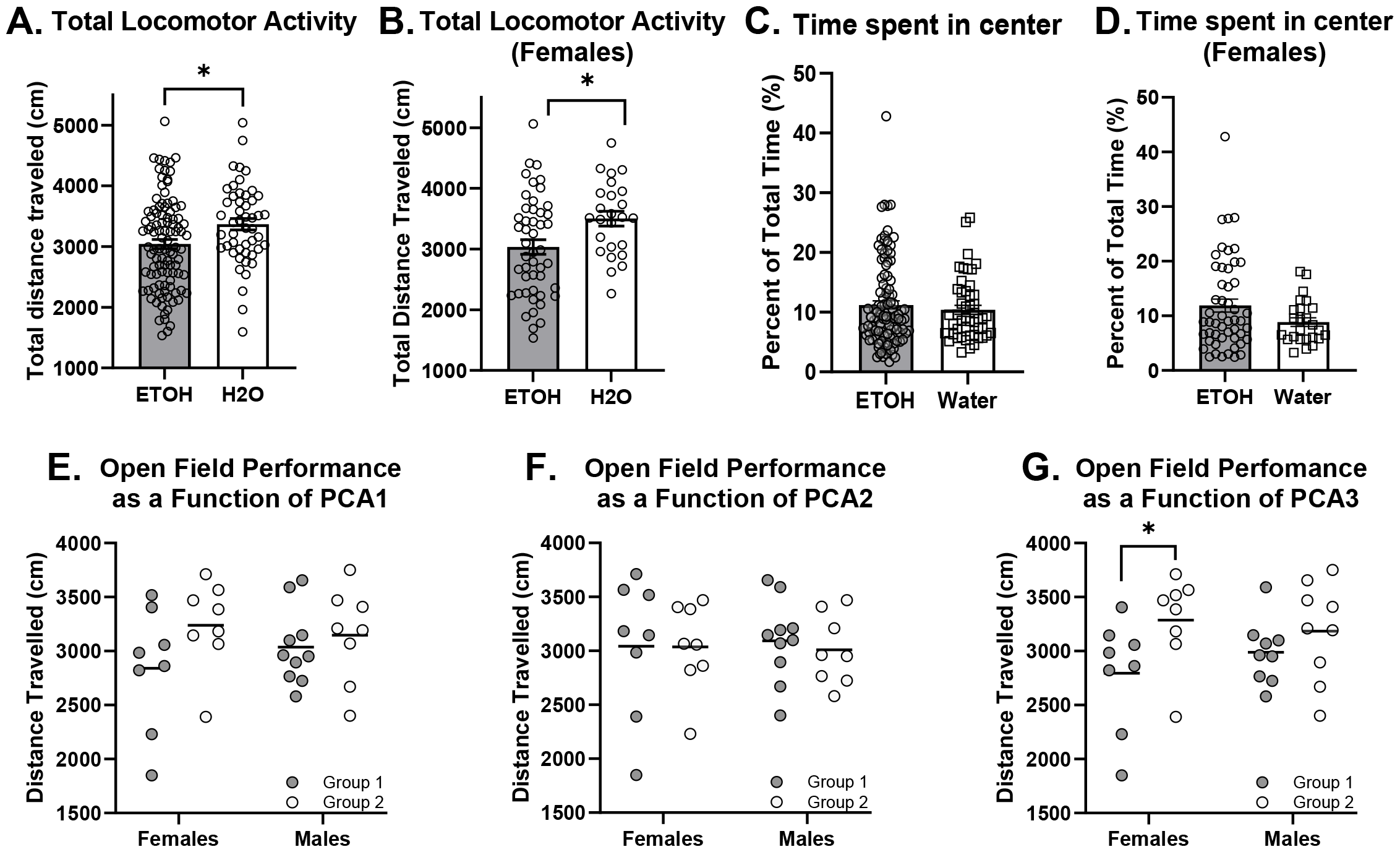
A) Total locomotor activity was significantly lower in ETOH exposed offspring (p = 0.0101). B) Treatment differences in total locomotor activity were driven by females (p = 0.0291). C) No significant treatment effect in percent of time spent in the center of the box. D) Percent of time spent in the center of the box for females. E) No significant correlation between OF performance and PCA1 (Females; p = 0.0674)(Males; p = 0.8118). F) No significant correlation with OF and PCA2 (Females; p = 0.8583)(Males; p = 0.6968). G) A significant correlation was found between OF performance and PCA3 among female mice (p = 0.0347) but not among male mice (p = 0.7713).

Next, we compared offspring behavior in the open field with our PCA defined MDPs. No relationship between PCA1 groups and OF behavior was found (Fig. 6E) [(Females; t(14) = 1.983, p = 0.0674)(Males; t(16) = 0.2421, p = 0.8118)]. Likewise, no correlation was found between OF behavior and PCA2 groups (Fig. 6F) [(Females; t(14) = 0.03307, p = 0.8583)(Males; t(54) = 0.3917, p = 0.6989)]. However, a significant relationship between total distance traveled in the OF and PCA3 MDPs was discovered in females [t(14) = 2.339, p = 0.0347] but not males [t(16) = 0.2956, p =0.7713] (Fig. 6G). No significant relationship between average cumulative maternal alcohol consumption and offspring open field performance was found for either sex (Females: p = 0.5018; Males: p = 0.1490).

### Allodynia

Differences in development of allodynia were investigated at mean postnatal day 164. Under baseline conditions (BL), light touch sensory thresholds demonstrated no significant differences between PAE and control mice. However, following a minor CCI, robust increases in hindpaw sensitivity were seen in female mice by post-surgery day 7 (Fig. 7A) with a main effect of treatment [F(1, 20) = 9.910, p = 0.0051] and a day by treatment interaction [F(6, 120) = 6.912, p<0.0001]. By post-surgery day 4, a main effect of treatment in males (Fig. 7B) [F(1, 12) = 53.38, p<0.0001] as well as a day by treatment interaction [F(6, 72) = 11.57, p<0.0001] was observed. The stimuli response in both sexes remained significant until termination of the study at 14 days post-surgery. Allodynia did not develop among the CCI or sham-injured control (water) mice. Due to the high PAE sample size necessary to evaluate offspring behavior using PCA derived MDPs, we were unable to perform this analysis with the mice underwent allodynia study.

**Figure 7.**
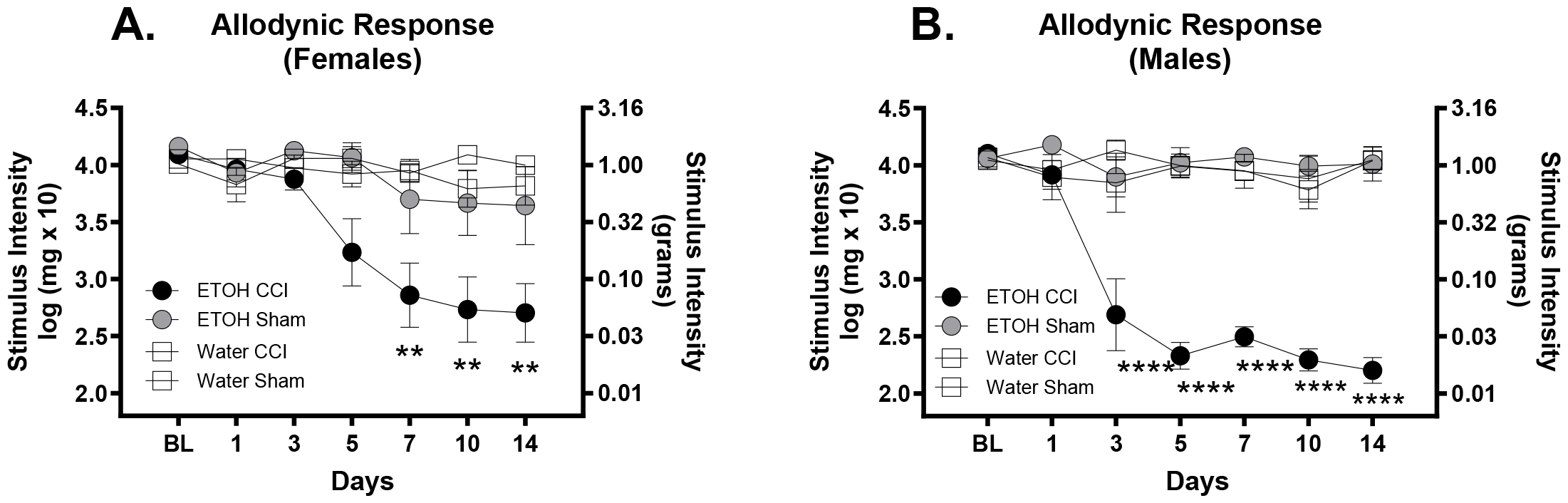
A) Females developed a main effect of treatment by day 7 (p = 0.0051). B) Males developed a main effect of treatment by day 5 (p<0.0001).

## Discussion

The results of this work replicate previous reports from our group that binge-like prenatal alcohol exposure leads to sex-specific alterations in behavior^1,2,5,16,17,24^ and extend them by identifying the *pattern* of gestational alcohol consumption as a critical intermediary attribute. Specifically, we found superior motor coordination on the rotarod in PAE males attributable to delayed weight gains, hypoactivity in a novel open field in PAE females, and sensitization to the development of allodynia in PAE mice which emerged earlier in males versus females. Importantly, we also identified two gestational alcohol-drinking patterns that were uniquely associated with alcohol frontloading and that differentiated neurobehavioral outcomes in exposed offspring. To our knowledge, this is the first report to link specific patterns of gestational binge drinking to behavioral outcomes in PAE offspring.

Using the VDM system together with daily 2-hour access to 20% unsweetened alcohol throughout gestation, we observed similar alcohol intake (Fig. 1A) and BEC profiles (Fig. 1B) to those we have previously reported using traditional sipper tubes with daily 4-hour access to 10% alcohol with saccharine ^14,16 1,25^. For example, Brady et al. (2012) reported almost exactly double the alcohol intake we report here, but this was expected given the amount of time to consume alcohol was also doubled (7.17 g/kg vs. 3.45 g/kg). BECs were approximately half of what we observed here (68.5 vs. 116.21 mg/dL), which we believe is attributable to an interaction between alcohol frontloading, and the longer time interval between alcohol consumption and blood sampling. That is, to the extent that peak BECs were similar between studies in both magnitude and timing (i.e. early in session due to frontloading), we would expected 4-hour BEC measurements to be approximately half those measured at the 2 hour time point, which they indeed were. Given these observations, it is likely that gestational alcohol exposure as experienced by the fetus is similar between these models.

Nonetheless, the model used here has several advantages. First, the extremely reliable linear relationship between ethanol intake and BECs in the 2-hour/20% paradigm provides more accurate estimations of BECs throughout gestation (Fig. 1C). Using this fact to our advantage, we discovered that while mean group-level alcohol consumption remained relatively stable, there was marked variability both between subjects on a given day and between days within a given subject (Fig. 1D-E). This observation emphasizes the fact that group means do not accurately reflect the experienced patterns of daily gestational alcohol exposure for a given litter. The removal of saccharine from all drinking solutions is a second advantage, in particular given the generally larger volumes of fluid consumed by control groups, which leads to differences in total gestational saccharine exposure. A final advantage, though obvious, lies in the ability of the VDM to accurately capture daily drinking patterns with high temporal resolution.

We leveraged the full power of the VDM system and observed drinking patterns consistent with our previous reports^14–16^, in which pregnant females reliably developed alcohol front-loading (Fig. 2A-B). Notably however, front-loading increased further over the course of gestation, particularly the last five days where pup development was most rapid (Fig. 2B). Assuming the increased gestational pup weight gain during this period leads to decreased distribution rate of alcohol to the central nervous system of the dam, this observation may represent compensatory drinking driven by the need to increase consumption rate to achieve similar BECs. Alternatively, increases could have been mediated by increased alcohol clearance at this stage in pregnancy, driven by increases in Aldehyde dehydrogenase (ALDH) and Alcohol Dehydrogenase (ADH4) enzymes^26^. Regardless of the reason, these data suggest that frequent alcohol consuming individuals who become pregnant may increase their alcohol consumption rate at later stages of gestation, an observation of clinical relevance that to our knowledge has not been studied in the humans. Notably, this late stage increase in frontloading may be related in part to an increase in motivation for alcohol developed during the 2 week alcohol drinking acclimation period prior to breeding, as frontloading is known to develop progressively over the initial 2 weeks of drinking^15,27^.

Building on these findings, we then used PCA to assess the most prominent patterns of alcohol consumption. We focused first on evaluating variation in drinking patterns within a single day (GD15). The two most dominant patterns of alcohol consumption on GD15 were differentiated from one another only following frontloading (Fig. 3B). Following frontloading, persistent drinkers (Group 1) continued a steady albeit slower rate of alcohol consumption, whereas Group 2 almost completely stopped drinking alcohol until approximately 30 minutes remained in the session. These drinking patterns were associated with significant differences in total intake and BECs (Fig. 3A; inset).

We then evaluated all gestational days together, restricting our initial PCA analysis to the first fifteen minutes of each session, followed by evaluation of the patterns of consumption over the entire 2-hour session. Group differences in alcohol drinking during the first 15 minutes (Fig. 3C) did not materialize into differences in total fluid consumption (Fig. 3D). However, they do reflect meaningful group differences in the rate of frontloading that would likely lead to differences in the rate of fetal alcohol exposure, and they *did* lead to differences in offspring development discussed below. Analysis of the entire 2-hr session (Fig 3E) revealed MDPs that were dissociated principally by the duration of frontloading (Fig 3F). Specifically, an approximately 1g/kg difference in alcohol consumption emerged around the first 15 minutes, and persisted to the end of the session. Collectively, these findings identify prominent MDP characteristics and set the stage for determining those most impactful to offspring behavior.

To do so, we first evaluated the impact of alcohol consumption on pup numbers and weight at P10 (Fig. 4B-C). Though persistent weight gain deficits have been attributed to children of heavy alcohol consumers, these effects are usually evident from birth^23,28^ and our study detected no weight differences in this early stage of development. However, by P57 when we began the first behavioral assessment, PAE males displayed significantly lower body weights (Fig. 4D). Our initial work using 20% ETOH solutions (but not 10%) also demonstrated weight difference starting only later in development (P21), but they were absent by P94^16^. These may be attributable to less food consumption in the PAE group, or, similar food intakes but differences in nutritional utilization. How and why weight differences were not observed earlier or later in development is not immediately clear.

Our assessment of motor coordination on the accelerating rotarod revealed significant differences in male mice only, with PAE outperforming their control counterparts (Fig. 5A). This initially curious finding led us to speculate that decreased body weight may have been driving differences in perceived motor performance. Indeed, we found a significant linear relationship between rotarod performance and body weight (Fig. 5B), and subsequently found a publication that corroborated this relationship in non-alcohol-exposed subjects^29^. These data replicate previous work by our Center using a third trimester model of alcohol exposure^17^, and provide important context to this and other work in the field on the relationship between development and perceived motor abilities. The increase in late stage gestational frontloading we observed in the current studies combined with superior motor abilities in a 3^rd^ trimester exposure model may indicate that high BECs towards later stages of pregnancy are the most impactful for future offspring rotarod performance. In support of this view, we found that the MDP associated with the first fifteen minutes of alcohol access dissociated male offspring into groups that differed in their performance on the rotarod (Fig. 5D). Specifically, Group 1 dams, which expressed an increased rate of frontloading, produced offspring who displayed better performance on the rotarod. Collectively, these data suggest that alcohol front-loading throughout gestation may be responsible for delays in weight gain among male mice, which paradoxically led to improved rotarod performance due to the known relationship between body size and task difficulty^29^, which we also observed (Fig. 5B).

Consistent with previous reports using similar DID models, we also found decreased overall locomotor activity among ETOH females in the OF^16,28^ (Fig. 6B). There was no evidence that these differences were due to differences in anxiety as measured by time spent in the center of the OF (Fig. 6C-D). However, decreased voluntary locomotor activity in the OF is potentially indicative of depressive or anxiety-like states^30^, so we cannot rule out this possibility. Regardless, we found that our 2-hour MDPs appeared to drive this effect. Specifically, Group 1 expressed more intense alcohol frontloading and a higher total alcohol consumption compared to Group 2, and their offspring travelled significantly shorter distances in the OF compared to their Group 2 counterparts. The other MDPs were not associated with the OF results, and total gestational alcohol consumed was not a significant predictor of OF performance. Together, these data indicate an intricate, sex-specific relationship between MDP characteristics and offspring behavior on the rotarod and OF. Future studies will be needed to further assess the contribution of between-session variation in these characteristics to offspring behavior, as the gestational timing of increases or decreases in particular drinking behavior may differentially contribute to neurodevelopment of the offspring.

Lastly, the most striking behavioral consequence of PAE was the induction of robust allodynia (Fig. 7A-B). This replicates our previous findings in male PAE rats^5,20,31,32^, and female PAE mice^21^, and extends them to mice of both sexes in our revised DID PAE paradigm. Importantly, male ETOH-exposed offspring were more sensitive to the development of allodynia, showing significant allodynic response earlier than females (post-operative day 5 vs 7). These results add to a growing body of literature indicating alterations in pain perception with PAE. Due to the high animal numbers required to evaluate relationships between MDPs and offspring behavior, we were unable to assess the potential contribution of MDPs to development of allodynia in these studies. However, our group is continuing to collect maternal drinking and offspring allodynia data in order to conduct such analysis in the future.

## Conclusion

The results presented here are the first to directly identify patterns of binge alcohol consumption leading to differential and sex-specific neurobehavioral effects in exposed offspring. Our data indicate that binge-level alcohol exposure *in utero* leads to allodynia in both sexes, weight and therefore motor function alterations in males, and decreases in exploratory activity in females. MDPs were found to be associated with these rotarod and open field assessments, in particular with alcohol frontloading – a drinking phenotype we were surprised to see increase further on the final days of gestation. These findings emphasize the importance of gestational alcohol MDPs, particularly the rate of alcohol consumption and how it may be influenced by alcohol availability, and further emphasize the need for future studies evaluating the impact of alcohol MDPs on offspring development. Additionally, we have demonstrated the relevance of the VDM system to studies of PAE and confirmed its use produces similar findings to those our group has obtained using older methods involving the daily weighing of individual bottles. The protocol outlined here therefore represents a major advancement in our first and second trimester-equivalent ETOH exposure model, which we will continue to leverage in our pursuit of mechanisms that can be leveraged as therapeutics.

## References

1. Brady, M. L., Allan, A. M. & Caldwell, K. K. A Limited Access Mouse Model of Prenatal Alcohol Exposure that Produces Long-Lasting Deficits in Hippocampal-Dependent Learning and Memory. Alcohol. Clin. Exp. Res. 36, 457–466 (2012).

2. Kenton, J. A., Castillo, V., Kehrer, P. & Brigman, J. L. Moderate Prenatal Alcohol Exposure Impairs Visual-Spatial Discrimination in a Sex-Specific Manner: Effects of Testing Order and Difficulty on Learning Performance. Alcohol. Clin. Exp. Res. 44, 2008–2018 (2020).

3. Hendricks, G. et al. Prenatal alcohol exposure is associated with early motor, but not language development in a South African cohort. Acta Neuropsychiatr. 32, 145–152.

4. Easey, K. E., Dyer, M. L., Timpson, N. J. & Munafò, M. R. Prenatal alcohol exposure and offspring mental health: A systematic review. Drug Alcohol Depend. 197, 344–353 (2019).

5. Sanchez, J. J., Noor, S., Davies, S., Savage, D. & Milligan, E.D. Prenatal alcohol exposure is a risk factor for adult neuropathic pain via aberrant neuroimmune function. J. Neuroinflammation 14, 254 (2017).

6. Clarke, M. E. & Gibbard, W.B. Overview of Fetal Alcohol Spectrum Disorders for Mental Health Professionals. Can. Child Adolesc. Psychiatry Rev. 12, 57–63 (2003).

7. Gosdin, L. K., Deputy, N. P., Kim, S. Y., Dang, E. P. & Denny, C.H. Alcohol Consumption and Binge Drinking During Pregnancy Among Adults Aged 18–49 Years — United States, 2018–2020. Morb. Mortal. Wkly. Rep. 71, 10–13 (2022).

8. Denny, C. H., Acero, C. S., Terplan, M. & Kim, S.Y. Trends in Alcohol Use Among Pregnant Women in the U.S., 2011–2018. Am. J. Prev. Med. 59, 768–769 (2020).

9. Muggli, E. et al. “Did you ever drink more?” A detailed description of pregnant women’s drinking patterns. BMC Public Health 16, 683 (2016).

10. Ikonomidou, C. et al. Ethanol-induced apoptotic neurodegeneration and fetal alcohol syndrome. Science 287, 1056–1060 (2000).

11. Lipinski, R. J. et al. Ethanol-Induced Face-Brain Dysmorphology Patterns Are Correlative and Exposure-Stage Dependent. PLoS ONE 7, e43067 (2012).

12. Bonthius, D. J. & West, J.R. Alcohol-induced neuronal loss in developing rats: increased brain damage with binge exposure. Alcohol. Clin. Exp. Res. 14, 107–118 (1990).

13. Kelly, S. J., Pierce, D. R. & West, J.R. Microencephaly and hyperactivity in adult rats can be induced by neonatal exposure to high blood alcohol concentrations. Exp. Neurol. 96, 580–593 (1987).

14. Maphis, N. M., Huffman, R. T. & Linsenbardt, D.N. The development, but not expression, of alcohol front-loading in C57BL/6J mice maintained on LabDiet 5001 is abolished by maintenance on Teklad 2920x rodent diet. Alcohol. Clin. Exp. Res. 46, 1321–1330 (2022).

15. Linsenbardt, D. N. & Boehm, S.L. Alterations in the Rate of Binge Ethanol Consumption: Implications for Preclinical Studies in Mice. Addict. Biol. 19, 812–825 (2014).

16. Boehm II, S. L. et al. Using drinking in the dark to model prenatal binge-like exposure to ethanol in C57BL/6J mice. Dev. Psychobiol. 50, 566–578 (2008).

17. Jacquez, B., Choi, H., Bird, C. W., Linsenbardt, D. N. & Valenzuela, C.F. Characterization of motor function in mice developmentally exposed to ethanol using the Catwalk system: Comparison with the triple horizontal bar and rotarod tests. Behav. Brain Res. 396, 112885 (2021).

18. Brigman, J. L., Mathur, P., Lu, L., Williams, R. W. & Holmes, A.Genetic relationship between anxiety- and fear -related behaviors in BXD recombinant inbred mice. Behav. Pharmacol. 20, 204–209 (2009).

19. Bennett, G. J. & Xie, Y.-K. A peripheral mononeuropathy in rat that produces disorders of pain sensation like those seen in man. Pain 33, 87–107 (1988).

20. Noor, S. et al. Prenatal alcohol exposure dysregulates spinal and circulating immune cell circular RNA expression in adult female rats with chronic sciatic neuropathy. Front. Neurosci. 17, 1180308 (2023).

21. Vanderwall, A. G. et al. Effects of spinal non-viral interleukin-10 gene therapy formulated with d-mannose in neuropathic interleukin-10 deficient mice: Behavioral characterization, mRNA and protein analysis in pain relevant tissues. Brain. Behav. Immun. 69, 91–112 (2018).

22. Milligan, E. D. et al. Thermal hyperalgesia and mechanical allodynia produced by intrathecal administration of the human immunodeficiency virus-1 (HIV-1) envelope glycoprotein, gp120. Brain Res. 861, 105–116 (2000).

23. Carter, R. C., Jacobson, J. L., Sokol, R. J., Avison, M. J. & Jacobson, S.W. Fetal alcohol-related growth restriction from birth through young adulthood and moderating effects of maternal prepregnancy weight. Alcohol. Clin. Exp. Res. 37, 452–462 (2013).

24. Licheri, V., Chandrasekaran, J., Bird, C. W., Valenzuela, C. F. & Brigman, J.L. Sex-specific effect of prenatal alcohol exposure on N-Methyl-D-Aspartate Receptor function in orbitofrontal cortex pyramidal neurons of mice. Alcohol. Clin. Exp. Res. 45, 1994–2005 (2021).

25. Brady, M. L. et al. Moderate Prenatal Alcohol Exposure Reduces Plasticity and Alters NMDA Receptor Subunit Composition in the Dentate Gyrus. J. Neurosci. 33, 1062–1067 (2013).

26. Badger, T. M., Hidestrand, M., Shankar, K., McGuinn, W. D. & Ronis, M.J. The effects of pregnancy on ethanol clearance. Life Sci. 77, 2111–2126 (2005).

27. Ardinger, C. E., Grahame, N. J., Lapish, C. C. & Linsenbardt, D.N. High Alcohol Preferring Mice Show Reaction to Loss of Ethanol Reward Following Repeated Binge Drinking. Alcohol. Clin. Exp. Res. 44, 1717–1727 (2020).

28. Kleiber, M. L., Wright, E. & Singh, S.M. Maternal voluntary drinking in C57BL/6J mice: advancing a model for fetal alcohol spectrum disorders. Behav. Brain Res. 223, 376–387 (2011).

29. Mao, J.-H. et al. Identification of genetic factors that modify motor performance and body weight using Collaborative Cross mice. Sci. Rep. 5, 16247 (2015).

30. Strekalova, T., Spanagel, R., Bartsch, D., Henn, F. A. & Gass, P.Stress-Induced Anhedonia in Mice is Associated with Deficits in Forced Swimming and Exploration. Neuropsychopharmacology 29, 2007–2017 (2004).

31. Noor, S. et al. Prenatal alcohol exposure potentiates chronic neuropathic pain, spinal glial and immune cell activation and alters sciatic nerve and DRG cytokine levels. Brain. Behav. Immun. 61, 80–95 (2017).

32. Noor, S. et al. The LFA-1 antagonist BIRT377 reverses neuropathic pain in prenatal alcohol-exposed female rats via actions on peripheral and central neuroimmune function in discrete pain-relevant tissue regions. Brain. Behav. Immun. 87, 339–358 (2020).

